# Targeted volume Correlative Light and Electron Microscopy of an environmental marine microorganism

**DOI:** 10.1101/2023.01.27.525698

**Authors:** Karel Mocaer, Giulia Mizzon, Manuel Gunkel, Aliaksandr Halavatyi, Anna Steyer, Viola Oorschot, Martin Schorb, Charlotte Le Kieffre, Daniel P. Yee, Fabien Chevalier, Benoit Gallet, Johan Decelle, Yannick Schwab, Paolo Ronchi

## Abstract

Photosynthetic microalgae are responsible for an important fraction of CO_2_ fixation and O_2_ production on Earth. Three-dimensional ultrastructural characterization of these organisms in their natural environment can contribute to a deeper understanding of their cell biology. However, the low throughput of volume electron microscopy (vEM) methods, along with the complexity and heterogeneity of environmental samples, pose great technical challenges. In the present study, we used a workflow based on a specific EM sample preparation, compatible with both light and vEM imaging in order to target one cell among a complex natural community. This method revealed the 3D subcellular landscape of a photosynthetic dinoflagellate with quantitative characterization of multiple organelles. We could show that this cell contains a single convoluted chloroplast and the arrangement of the flagellar apparatus with its associated photosensitive elements. Moreover, we observed chromatin features that could shed light on how transcriptional activity takes place in organisms where chromosomes are permanently condensed. Together with providing insights in dinoflagellates biology, this proof-of-principle study illustrates an efficient tool for the targeted ultrastructural analysis of environmental microorganisms in heterogeneous mixes.

## Introduction

Electron microscopy (EM) has played an essential role in understanding cell biology by revealing the intracellular organization in a vast variety of cells, tissues and small organisms (Introduction to Electron Microscopy of Cells, 2007). More recently, vEM methods have given the possibility to visualize ultrastructure in 3D (Peddie et al., 2022), opening the way to new discoveries. However, to date, these methods have been rarely applied to study marine microplankton. Microplankton are microorganisms that populate aquatic ecosystems. They include a wide variety of species ranging from prokaryotes to eukaryotic microalgae. Our understanding of their cell biology is still very limited. Indeed, working with highly heterogeneous environmental samples is very challenging, and only a small fraction of species can be cultured in the laboratory (Dixon and Syrett, 1988; Oliveira et al., 2020). Therefore, new methods to characterize these cells in their native ecosystem is highly needed.

In the present study we present a workflow to characterize by vEM a microorganism from environmental samples addressing the bottlenecks discussed previously. For this proof of principle, we focused on a dinoflagellate cell. Dinoflagellates represent a considerable fraction of plankton and play an important role in the aquatic food chain (Vargas et al., 2015). There are close to 2400 described species, which are highly heterogeneous in morphology, trophic mode and distribution (Gomez, 2012). Approximately half of them are primary producers contributing to O_2_ production and CO_2_ fixation on the planet (Gomez, 2012). As most marine planktonic cells, dinoflagellates are difficult to maintain in culture (Dixon and Syrett, 1988). Therefore, only a small fraction of species has been thoroughly investigated. Nonetheless, a coarse picture of the subcellular characteristics of dinoflagellates can be extracted from past EM studies conducted on cultured species.

One of the most striking features of these organisms is that they have numerous (up to 200) chromosomes (Bhaud et al., 2000) which stay permanently condensed throughout their cell cycle (Gautier et al., 1986). A fraction of these organisms possesses rigid cellulose plates located in a single layer of flattened vesicles lying under the plasma membrane, forming the theca. This contributes to the distinctive shape of the cells and is often used for taxonomic classification. Characteristic organelles of the dinoflagellate are trichocysts, described as rod shaped crystalline structures with squared cross-section profile (Bouck and Sweeney, 1966). These structures can be extruded from the cell (Westermann et al., 2015), potentially as a defense mechanism. However, their function is still highly debated (Plattner, 2017). Moreover, dinoflagellates generally present a Golgi complex hemispherically distributed above the nucleus (Dodge, 1971) as well as secretory organelles of various shapes and content named mucocysts (Hoppenrath, 2017). Dinoflagellates generally present two flagella, important for cellular movement (Dodge, 1971). Closely associated to the flagella, these organisms can have photosensitive structures, called eyespots, potentially responsible for directionality of their movement (Dodge, 1984). As the picture described above is derived from TEM studies, with a few exceptions where vEM was used (Decelle et al., 2021; Decelle et al., 2022; Gavelis et al., 2019; Uwizeye et al., 2021a; Uwizeye et al., 2021b), the 3D understanding of their organization is still largely lacking. Importantly, it has been described that some cells can lose specific structures, as for instance the eyespot, when kept in culture (Moldrup et al., 2013). Therefore, implementing culture independent methods to study planktonic cells in their native ecosystem is truly important to better understand the cellular biology of these ecologically relevant microorganisms.

Here we present a workflow that enables the identification of specific microalgal taxa of interest in a highly heterogenous environmental sample, containing hundreds of cells of diverse species. We used an EM sample preparation method that allows correlative analysis of the three-dimensional cell fluorescence pattern and focused ion beam – scanning electron microscopy (FIB-SEM) acquisition. To this aim, we have further optimized the workflow presented by Ronchi et al., 2021. This has enabled us to efficiently generate a vEM dataset of a specific environmental cell of interest, in this case a photosynthetic dinoflagellate. The analysis of the 3D ultrastructure allowed us to understand the intracellular organization of the organelles, revealing new insight in the biology of this microorganism.

## Results

### Sample preparation and cell identification

Preparation of environmental samples for ultrastructural studies by EM is highly complex. The quality of preservation is very time sensitive since planktonic microorganisms are delicate and susceptible to distortions before and during fixation (Truby, 1997). To help overcome these challenges, the samples were cryo-immobilized by high-pressure freezing within 2 hours after collection in a custom set up implemented in a marine station (see Material and Methods).

Each frozen sample contained hundreds of cells representing diverse plankton taxa. As most microalgae display characteristic autofluorescent properties, we decided to utilize light microscopy to target specific cells of interest. We therefore performed a freeze substitution method for preserving fluorescence by using a low amount of heavy metals and embedding in Lowicryl HM20 (Kukulski et al., 2011; Nixon et al., 2009; Porrati et al., 2019; Ronchi et al., 2021). Confocal 3D imaging of the resulting block revealed that the autofluorescence pattern was preserved after sample preparation and could be detected in the block for the entire thickness of the HPF material (200 µm) (Fig 1A, Video S1). Using excitation light at 488 nm and 633 nm along with transmitted light to visualize cell morphology, we could distinguish different families in the block (Fig. 1B-M), and identify damaged cells to exclude from downstream processing. As chlorophyll *a* had been reported to be autofluorescent in far red (Hense et al., 2008), we expected to be able to discriminate between photosynthetic and non-photosynthetic organisms by the presence of an emitted signal upon illumination at 633 nm.

**Figure 1.**
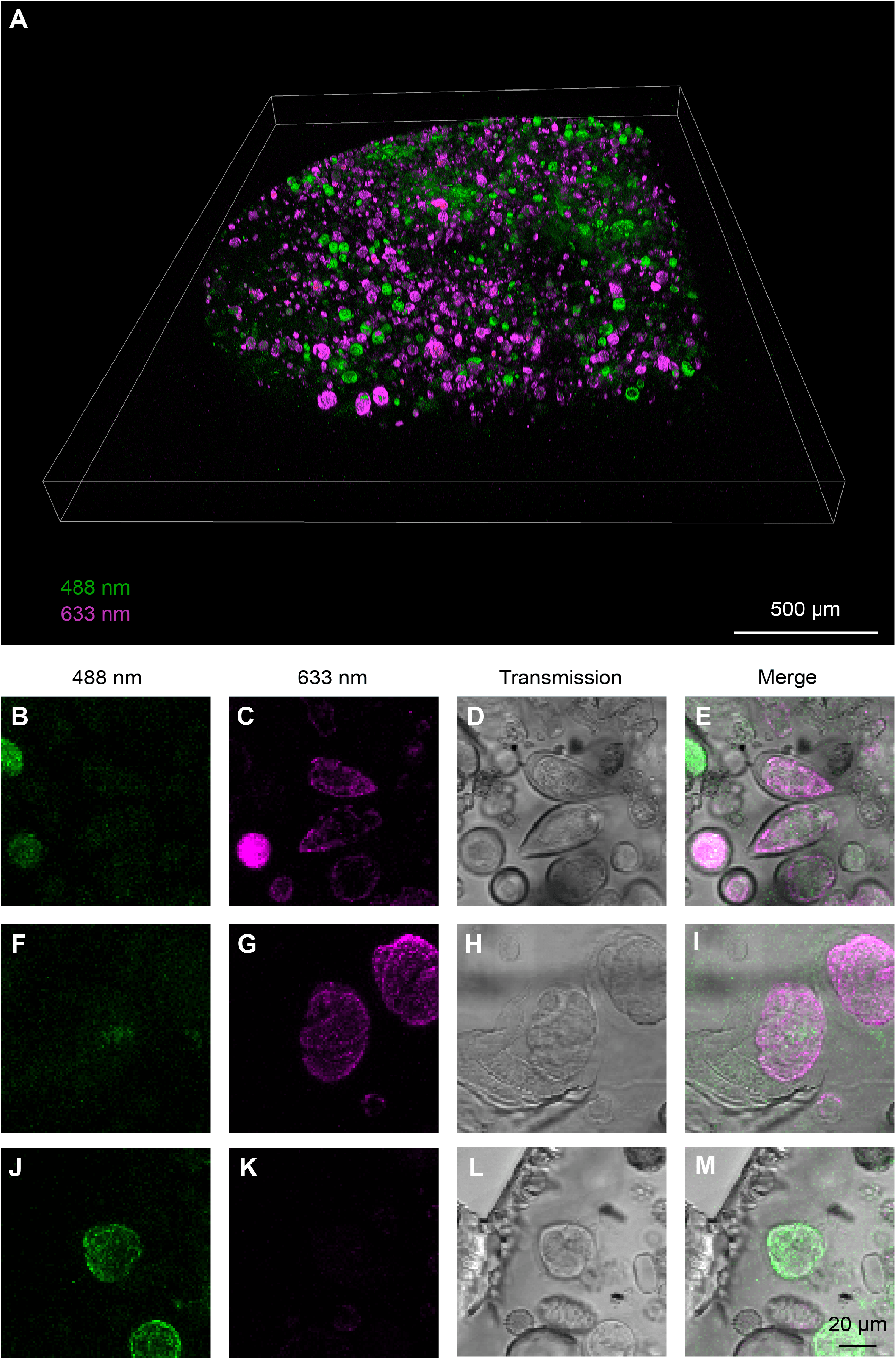
Confocal characterization of the HPF/FS planktonic sample embedded in a plastic block. A) 3D rendering of the 2-color tiled Z-stack confocal acquisition of the resin block. B-E; F-I; J-M) Fluorescence and transmitted light imaging of 3 different cells from A. The imaging settings are the same for the different cells in each channel. Maximum intensity projections of the confocal stacks are displayed for both fluorescence channels. For the transmitted light channel, single slices are shown. Fluorescence and transmitted light are overlaid in E,I,M. The cells were putatively identified as belonging to: B-E) genus *Prorocentrum*, F-I) infrafylum Dinoflagellata and J-M) genus *Protoperidinium*.

Considering the information provided by light microscopy, for this study we aimed for the vEM analysis by FIB-SEM of a plastid bearing dinoflagellate. Therefore, using transmitted light we selected a cell that presented a transversal groove and one longitudinal located antiapically (Video S1), as it is typical of many dinoflagellates. From the cells showing this particular shape, we further selected an organism that displayed far red signal when excited at 633 nm. The autofluorescence pattern consisted in a globular shape in the center of the cell and more defined patches on the cell periphery (Fig. 2A-D, Video S1), which we hypothesized to be chloroplasts.

**Figure 2.**
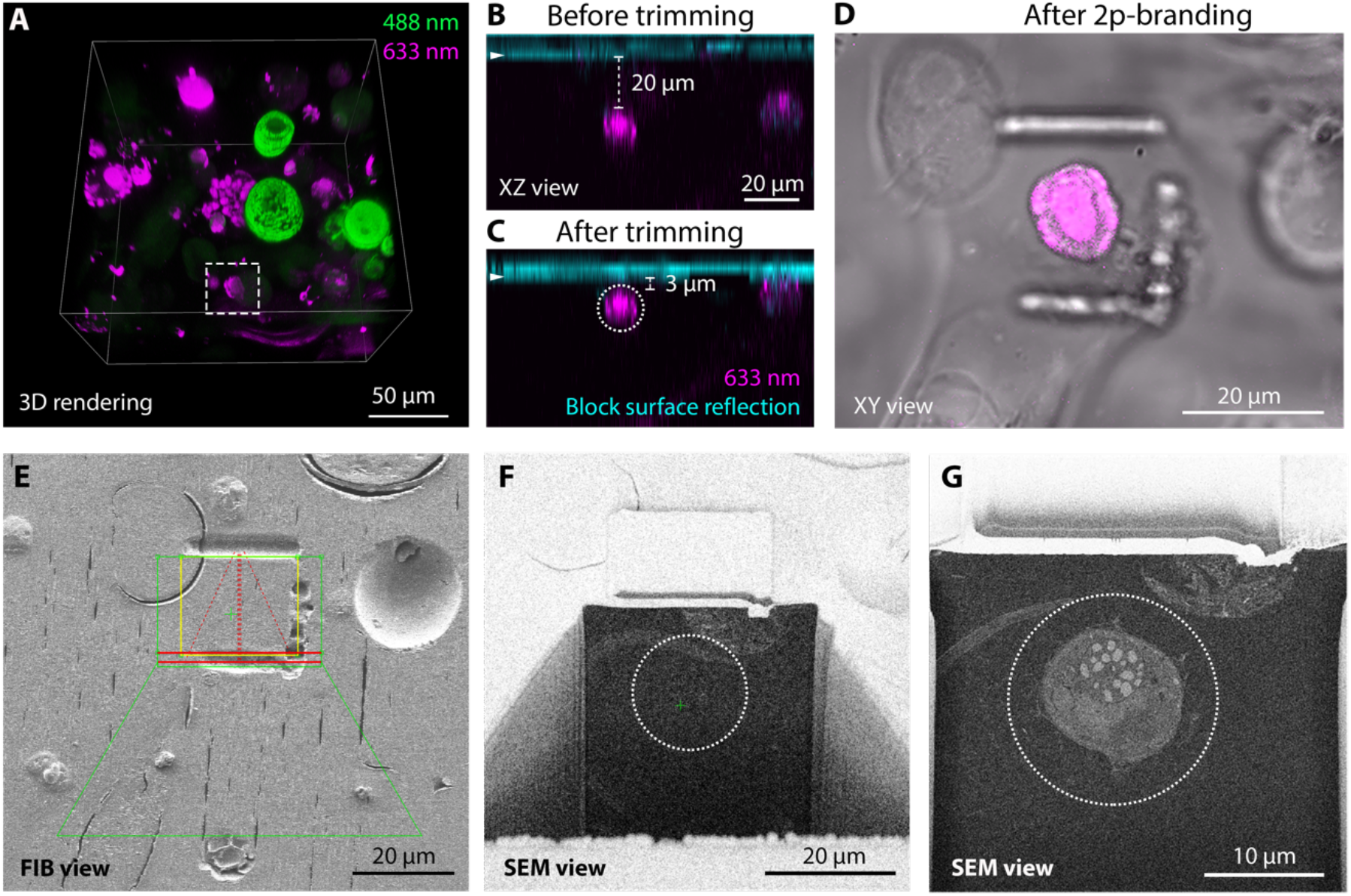
Targeting of a cell of interest (a photosynthetic dinoflagellate). A) 3D rendering of a high-resolution confocal stack in a selected area allows the identification of the target dinoflagellate cell (dashed square). B,C) Confocal XZ views of the cell of interest in the resin block before (B) and after (C) the trimming steps. The arrowhead indicates the block surface, as visualized in the reflection channel (cyan). Distance between the upper edge of the cell and the block surface is displayed. The dashed circle corresponds to the target position of the cell to be acquired by FIB-SEM. D) NIR branding of the block surface generates landmarks around the cell of interest, visualized by transmitted light. E) The embossed lines generated by the branding are visible by FIB imaging. These lines are used to define the region to be acquired by FIB-SEM. The overlaid profiles (green trapezoid and rectangle, red lines and yellow bounding box) illustrate the software (Atlas) sample preparation shapes used to define the slice-and-view acquisition. F) SEM view of the imaging surface after FIB sample preparation, right before starting the acquisition. The cell of interest is not exposed yet and the dashed circle represents its predicted position from Fig. 2C. G) Low magnification SEM overview (keyframe) during the acquisition, showing the precision of the ROI prediction. The dashed circle is in the same position as in F.

In order to image the cell of interest by FIB-SEM, we optimized the strategy reported by Ronchi et al., 2021, based on a 2-step targeting workflow. First, the distance of the cell of interest from the block surface was measured from a confocal stack (Fig. 2A,B). To determine the exact position of the block surface, we used the reflection of the laser light at the interface between materials with different refractive indexes (water used as a mounting medium and Lowicryl of the block, Fig 2B, arrowhead). Then we removed the measured thickness of resin present above the cell of interest with a trimming knife mounted on an ultramicrotome in three iterations of imaging and trimming. Once the cell was located just below the surface (Fig.2C), we branded landmarks on the block surface using a near infrared (NIR, “2-photon”) laser in order to later facilitate its targeting at the FIB-SEM (Fig. 2D). Indeed, such branded landmarks were easily identified at the SEM and were used to define the FIB milling area (Fig. 2E). A trench was then opened to approach the cell according to its predicted location (Fig. 2F). Using the measurements of the confocal stack (Fig. 2C, dashed circle), we could predict precisely the position of the acquisition window where the cell would appear (Fig. 2F, dashed circle). The FIB-SEM automated acquisition was then started and the entire cell was acquired at 8 nm isotropic voxel size (Fig. 2G, Video S2. The complete dataset is made available for download on EMPIAR). The complete dataset and associated segmentation can be visualized and interacted with using Mobie (https://github.com/mobie/environmental-dinoflagellate-vCLEM, see Material and methods, Video S1). The quality of the data acquired was consistent with previously published volume correlative light and electron microscopy datasets (Dimprima et al., 2022; Porrati et al., 2019; Ronchi et al., 2021).

By overlaying the light microscopy and the FIB-SEM data, we tried to confirm the nature of the autofluorescence signal. We observed that the far-red signal matched with the position of the chloroplast and the nucleus (Fig. 3A-C, Video S1). While we cannot explain the autofluorescence in the nucleus, the overlap with the plastid is justified by the presence of chlorophyll (Hense et al., 2008). This confirms that the far-red signal from microorganisms within the block can be used for the identification of photosynthetic species.

**Figure 3.**
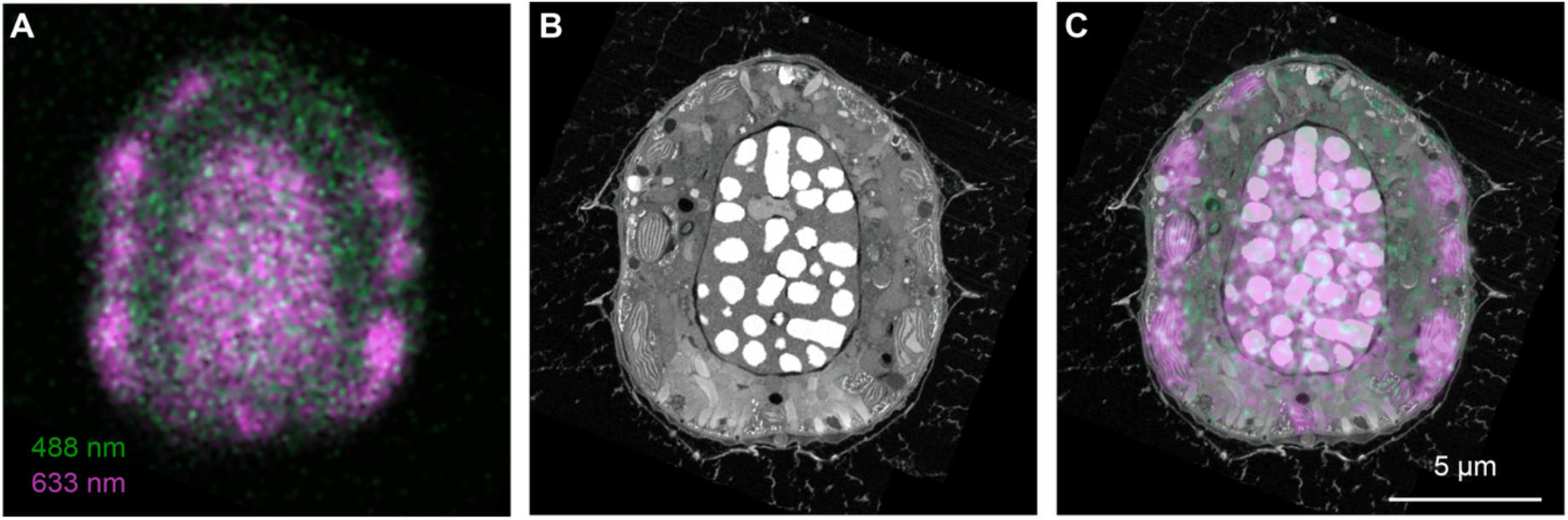
Overlay of autofluorescence signal and ultrastructure from vEM. A) Fluorescent pattern of the cell of interest. The image shows a confocal slice along the longitudinal axis, optical thickness of 2.2 µm. B) Single orthoslice through the FIB-SEM volume in the region corresponding to the fluorescence signal. C) Fluorescence and FIB-SEM overlay. A single slice of the overlay is shown. The pattern of the 633 nm excited signal clearly overlaps with the position of the chloroplast and nucleus.

### Ultrastructural characterization

We then segmented a set of characteristic organelles of the cell from the FIB-SEM stack (Fig. 4, Video S2) and performed subsequent morphometric analysis. The cell measured 16 µm in height and 13.5 µm in width, for a total volume of 1009 µm^3^. It showed a conical epitheca as well as a wide and deep cingular girdle. The cell surface was nicely preserved allowing most thecal plates to be counted and described individually (with the exception of the sulcal plates). Elucidation of the plate tabulation was important for taxonomical identification. Traditionally, this analysis is done using SEM, however we show here that it can be also obtained from the reconstruction of the FIB-SEM volume. From our analysis, we could observe the following thecal arrangement according to the Kofoidian system (Fensome, 1993) : x, 4’, 3a, 7’’, 4c+T, 5’’’, 2’’’’. The topography of the different thecal plates displayed circular pores surrounded by small knobs or bumps (Fig. 4A). The pores were either linearly arranged as, for instance, above and under the cingulum or distributed with various densities throughout a given plate. Small knobs were also distributed unevenly throughout the thecal plates. Given the size range, outer morphology and the tabulation, we identified this organism as *Ensiculifera tyrrhenica (*synonym to *Pentapharsodinium tyrrhenicum* (Balech)), a dinoflagellate belonging to the class Dinophyceae and order Peridiniales. Analysis of the environmental sample collected in parallel and processed for topography SEM confirmed the presence of this species (Fig. S1). Interestingly, the position of the thecal openings and details of the ornamentation of the organisms analyzed with the 2 methods were extremely similar (Fig. S1), validating our FIB-SEM imaging-based analysis as a tool for determining taxonomic features. However, vEM has the additional advantage to provide insights on the intracellular morphology of the cell, which cannot be appreciated with SEM only.

**Figure 4.**
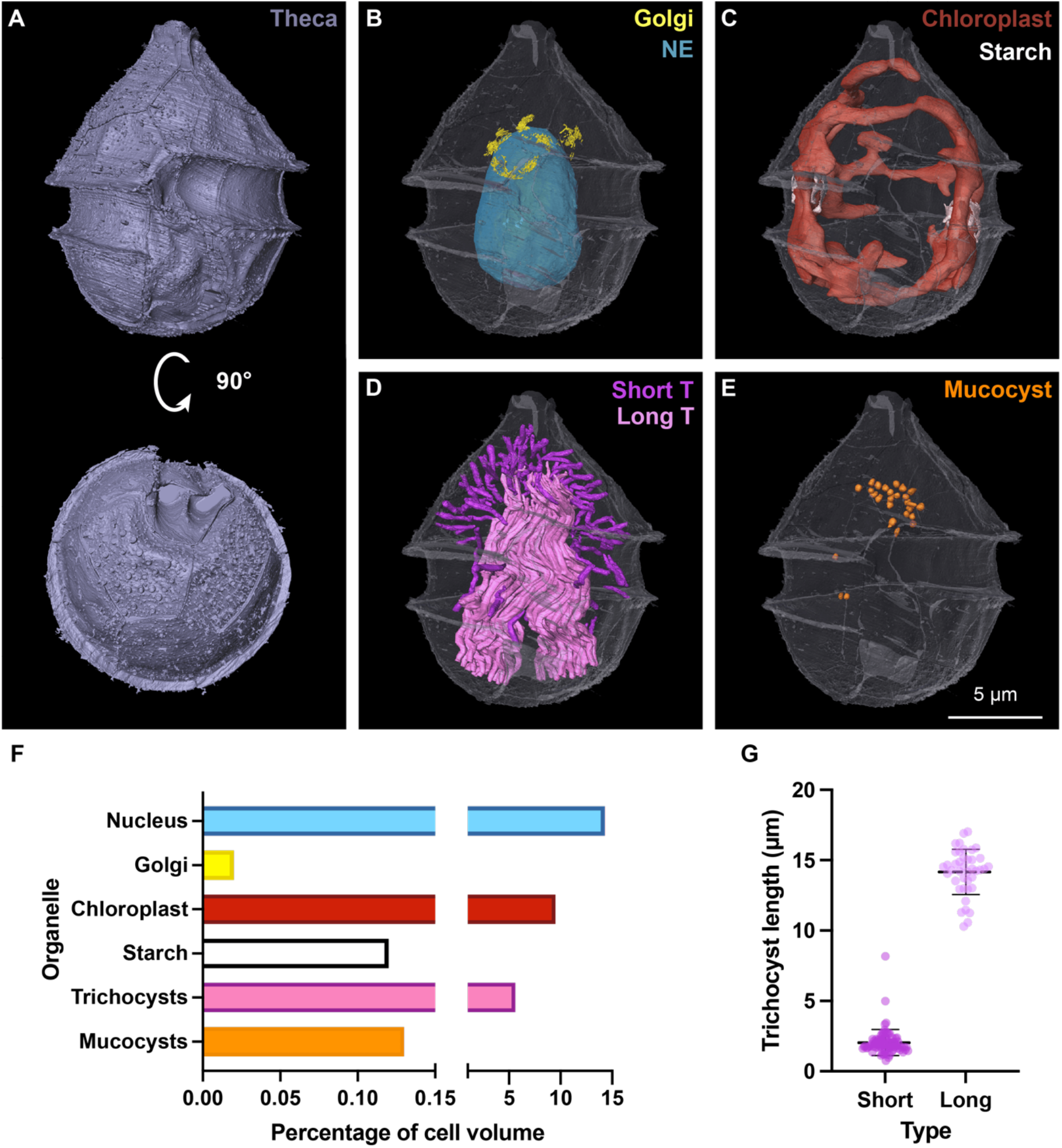
Morphometrics of organelles in the targeted photosynthetic dinoflagellate. A) 3D rendering of the theca in ventral (top) and antiapical views (bottom). The antiapical view in the lower panel shows the pore arrangement on the lower plates. All images of intracellular organelle segmentation (B-E) are shown in the same orientation as the top panel (ventral view), with the theca shown in transparency. B) Segmentation of the nucleus/nuclear envelope (NE) (cyan) and Golgi apparatus (yellow). C) Segmentation of the single convoluted chloroplast (red) and associated starch (white). D) Segmentation of the trichocysts. Short linear ones are shown in magenta. Long and convoluted ones in light pink. E) Segmentation of the mucocysts (orange). F) Volume of the segmented organelles, expressed as relative percentage of the full cell volume (1008.76 µm^3^). G) Size distribution of the two classes of trichocysts. N= 80 for the short and 41 for the long class.

Segmentation of some intracellular organelles allowed us to assess their position as well as size, shape and volume (Fig. 4, Video S2). The nucleus, representing 14.3% of the cell’s volume, has an elliptical shape and is located centrally in the posterior part of the cell, with its major axis aligned with the long axis of the cell (Fig. 4B,F). The Golgi apparatus is organized in 12 stacks located in the ventral apical region of the cell close to the nucleus (Fig. 4B, Fig. S2A-B), and occupied 0.02% of the cell volume (Fig. 4F). Classical taxonomical description of the genus *Ensiculifera* based on light microscopy reports the presence of reticulated chloroplasts (Li et al., 2020a). However, identification of the number of chloroplast or the pyrenoid distribution is difficult to interpret from light microscopy or even TEM images. Our vEM analysis allowed to unambiguously resolve the 3D organization of this organelle as a single convoluted and interconnected structure (Fig. 4C). The chloroplast represented 9.5% of the cell total volume (Fig. 4F). From the raw data, we could also appreciate the thylakoids and pyrenoid organization (Fig. S2C-D). Associated to the chloroplast, we observed the presence of starch, representing 0.12% of the cell volume and confined around two opposite lobes of the plastid where the pyrenoid was located (Fig. 4C,F, Fig. S2C-D). Of note, the sample was collected before sunrise, and a low amount of starch is compatible with night starch consumption (Seo and Fritz, 2002). We further analyzed the trichocysts, organelles typically found in dinoflagellates described as rod shaped crystalline structures originating from the Golgi area with a squared profile when cut transversally (Bouck and Sweeney, 1966). Our 3D analysis confirmed these general features of the trichocysts (Fig. S2E-F), but further allowed us to divide them in two classes, based on their length and distribution (Fig. 4D,G). One class comprises short (2.04 ± 0.92 µm, Fig. 4G) and straight structures, often perpendicular to the plasma membrane in the apical region (Fig. 4D, magenta). A second class formed a bundle of long (14.17 ± 1.61 µm, Fig. 4G), twisted and intricated trichocysts stretching along the longer cellular axis (Fig. 4D, light pink). Contrary to our expectations, neither class aligned with the thecal circular openings. Altogether, trichocysts occupy a significant fraction of the cell volume (5.61%, Fig. 4F). We further analyzed secretory organelles, described as mucocysts in dinoflagellates. We identified 30 amphora shaped structures (Fig. S2G-H), with an average volume of 0.046 ± 0.017 µm^3^. Altogether, mucocysts occupy 0.13% of the cell volume and are clustered under the plasma membrane in the posterior apical region of the cell (Fig. 4E,F).

Next, we performed a detailed characterization of the nuclear organization (Fig. 5). Chromatin segmentation revealed that the nucleus contained 105 condensed chromosomes (Fig. 5A,B). Their volume is on average 0.510 µm^3^ ± 0.162 µm^3^. Interestingly we could also observe two chromosomes located adjacent to the nucleolus which appeared smaller (respectively 0.025 µm^3^ and 0.004 µm^3^) compared to the other chromosomes. A detailed analysis of these structures showed threads of electron-dense material originating from the chromosomes and extending within the nucleolar space in a convoluted manner (Fig. 5C,D). The electron density properties, similar to those of neighboring chromosomes (Fig. 5A, arrowhead), suggest that these filamentous structures could be chromatin in an intermediate compaction state.

**Figure 5.**
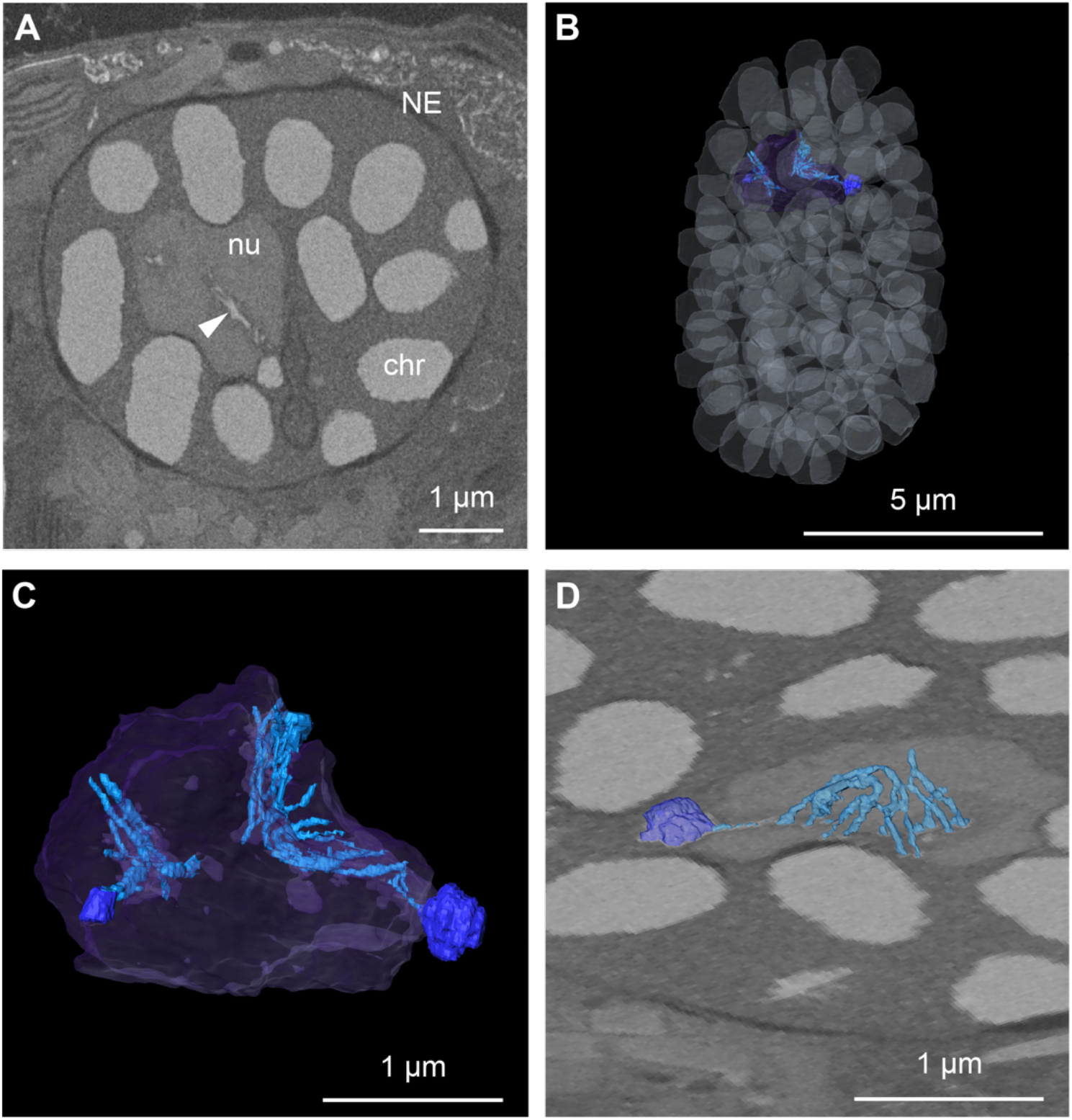
Nucleus and chromatin organization. A) Single orthoslice through the FIB-SEM volume in the nuclear region. (NE = nuclear envelope, nu = nucleolus, chr = condensed chromosome). The arrowhead highlights filamentous structures originating from a small chromosome and expanding in the nucleolus. B) 3D rendering of the segmentation of the chromatin (white in transparency), nucleolus (purple in transparency) and filamentous structure (light blue) associated with small chromosomes located adjacent to the nucleolus (dark blue). C) Close up view of the segmentation of nucleolus (purple in transparency) with associated small chromosomes (dark blue) and extended filamentous structure (light blue). D) Rendering of the segmentation of the intranucleolar filamentous chromatin structure overlaid with an image of the raw data, illustrating the connection between the filament and the small chromosome associated with the nucleolus.

Furthermore, we looked at the flagella, characteristic structures of dinoflagellates and at the associated eyespot (Fig. 6A), a putative photosensitive structure (Colley and Nilsson, 2016). The cell displayed 2 flagella, protruding from basal bodies located underneath the intersection between the sulcus and cingulum (Fig. 6B). These structures were elongated in the space between the plasma membrane and theca (Fig. 6C), a position that to our knowledge had not been described previously. The longitudinal flagellum appears very long and wrapping half of the cell perimeter, whereas the transverse flagellum is very short, as it may have been affected during sample collection. The eyespot could be visualized behind the sulcus groove (Fig 6B), as reported for other peridiniales (Dodge, 1984; Li et al., 2020b). The eyespot consisted here in a single layer of globules within the chloroplast and localized parallel to the longitudinal flagellum (Fig. 6E,F). This type of arrangement has been previously described for other organisms from the family of Peridiniaceae (Calado et al., 1999; Moestrup and Daugbjerg, 2007). For many dinoflagellate species the presence of flat arrays of laterally connected microtubules in the basal body area have been reported (Calado and Moestrup, 2002; Calado et al., 1999), which were suggested to play a role in light dependent movements (Dodge, 1984). Although the size of microtubules is at the limit of what can be resolved using our FIB-SEM imaging settings, we were able to visualize two structures resembling filamentous arrays, each one closely associated to a basal body (Fig, 6D-F). Taking advantage of the high density provided by the bundling of microtubules, we were able to localize them and determine their spatial arrangement. While the filaments related to the basal body of the longitudinal flagellum were directed toward the cell surface, following the curvature of the plasma membrane, the filaments related to the basal body of the transversal flagellum were directed towards the inner part of the cell (Fig. 6E,F).

**Figure 6.**
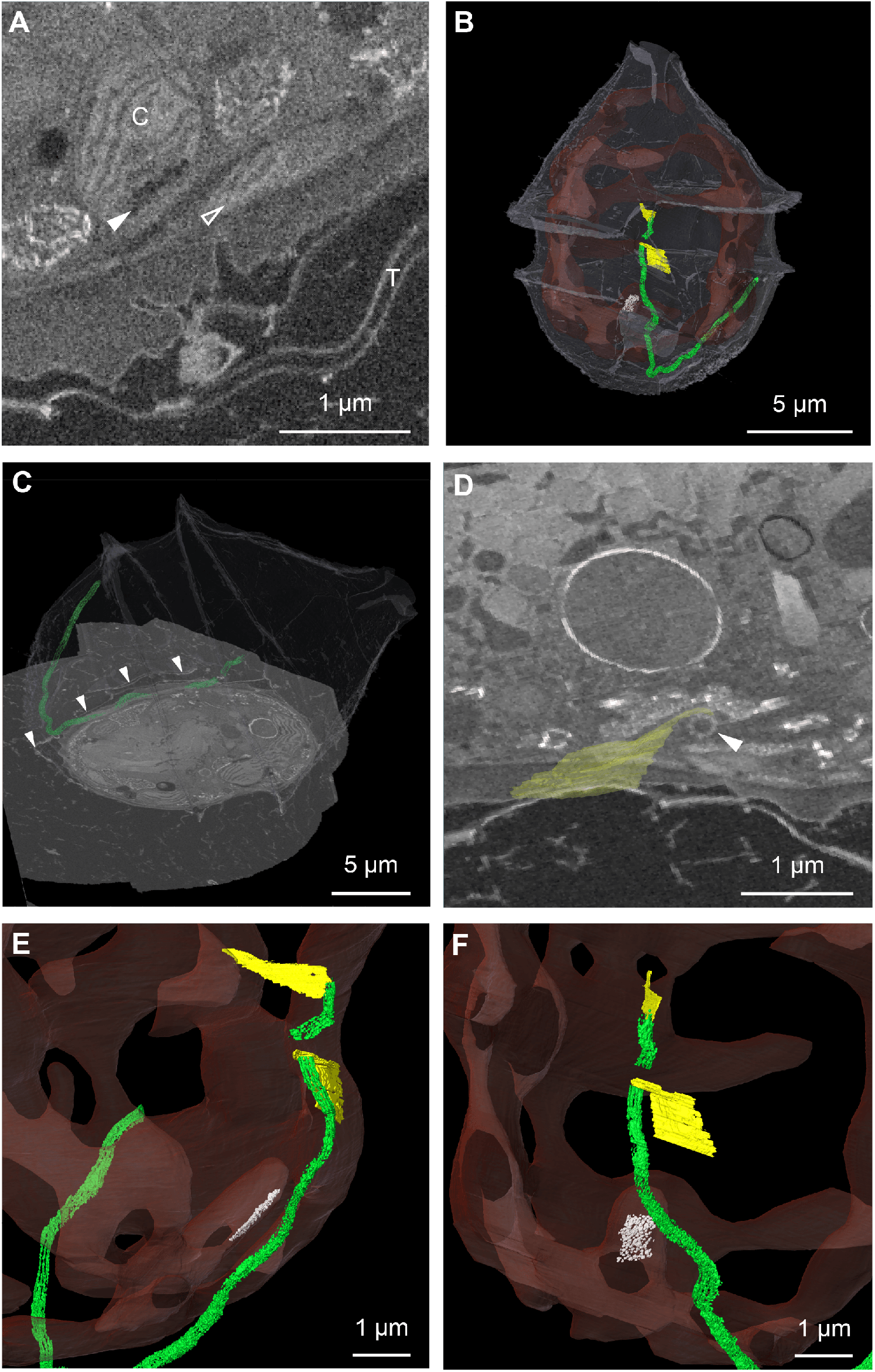
3D organization of the flagellar apparatus and eyespot. A) Single orthoslice through the FIB-SEM volume in the eyespot region. (C = chloroplast, T= theca, full arrowhead pointing towards the eyespot and outlined arrowhead pointing towards the longitudinal flagellum) B) 3D rendering of the segmentation of the eyespot (white), located within the chloroplast (red in transparency), of the flagella (green) and associated filaments (yellow). The theca is shown in white in transparency. C) Rendering of the segmentation of the longitudinal flagellum overlaid with a slice of the FIB-SEM volume. Arrowheads point at the position of the theca, demonstrating that the flagellum extends inside the theca. D) Rendering of the segmentation of a microtubule sheet (yellow) overlaid with a slice through the volume, showing the basal body of the flagellum (arrowhead). E-F) Close up on the segmentation of the chloroplast (red in transparency), flagella (green) and associated filaments (yellow), as well as the eyespot (white) located within the chloroplast.

## Discussion

In this study, we have presented a 3D correlative light and electron microscopy workflow for the identification and ultrastructural analysis of single organisms from a very heterogeneous environmental marine sample.

When using vEM, it is often necessary to limit the acquisition to a few “representative” individuals in a population because of the intrinsic low throughput of the method. In contrast to laboratory monocultures where inter-individual variability is relatively low, environmental samples can contain hundreds of species. Therefore, one of the biggest hurdles of meaningful 3D ultrastructural analysis of field samples is the identification and targeting of specific microorganisms. Using fluorescence profiling within the block, our method provides a novel way to explore heterogeneous samples to identify and select candidates for vEM acquisition. A similar workflow has been shown to work for exogenously expressed fluorescent proteins (Ronchi et al., 2021) or small molecules live dyes (Dimprima et al., 2022). We have now demonstrated that the same principle can be applied to endogenous fluorescence signals, opening the way for environmental sample analysis. From the shape and fluorescence pattern observed in the confocal stack, we were able to target a plastid-bearing unicellular armored dinoflagellate. Following FIB-SEM imaging we could confirm our prediction and identify the species by the thecal organization as *Ensiculifera tyrrhenica*.

Another advantage that comes with the precision of the workflow used is the reduction of the acquired FIB-SEM volume. Indeed, while a low precision targeting would lead to the acquisition of a large buffer volume around the region of interest, with this workflow we were able to restrict the acquisition very precisely around the cell of interest. This allowed us to optimize the imaging time and generate the entire dataset of the cell (∼15 µm diameter) in less than 48 h. In the future, using this dataset as a reference and taking advantage of specific autofluorescence signatures, it may be possible to further restrict the vEM acquisition to a subcellular volume to answer specific biological questions.

Our study is not only a methodological proof of concept but represents one of the first examples of vEM on environmental samples. More specifically, we describe here the subcellular organization of *Ensiculifera thyrrenica* for the first time. As datasets of this sort are rare and precious for the community, we believe it is important to make them available. Thus, the raw dataset as well as the segmentations described in this study are accessible in an easily browsable format using Fiji through the Mobie plugin (Pape et al., 2022).

With this vEM dataset, we could visualize the positioning of various structures which showed that the cell is highly polarized. Indeed, a subset of organelles are particularly concentrated in the apical region of the cell such as the Golgi apparatus and mucocysts. The distribution of the trichocysts, and in particular the short ones, appears polarized as well. They radiate from the Golgi area and are directed towards the apical plasma membrane suggesting they could be mature and ready for extrusion. Furthermore, the 3D analysis allowed us to observe the position of subcellular structures relative to one another as for the eyespot, flagella and associated filaments, which would be very difficult using other methods. The close association of the arrays of filaments with each basal body suggests that these microtubules could play a role in orienting the movement of the flagella. A higher resolution imaging of this area could further reveal whether these arrays are associated with the eyespot, which may in this way determine the directionality of the movement as suggested by Dodge, 1984.

The permanently condensed nature of dinoflagellate chromosomes (Gautier et al., 1986) raises questions concerning how they transcribe their genomes. Our high resolution 3D visualization of the nucleus allowed us to notice the presence of a filamentous structure originating from small chromosomes and extending inside the nucleolar volume. As the properties of these threads are very similar to the chromosomes they are originating from, we believe they represent a chromatin intermediate unfolding state. This arrangement is consistent with potential intranucleolar transcriptional activity which has been previously hypothesized (Géraud et al., 1991), but never demonstrated.

This work also highlights the importance of correlating 3D light and EM data beyond targeting purposes. The subcellular precision of such correlation allowed us to assign the emission of a fluorescent signal (far-red) to a specific organelle (chloroplast). Further studies, by mapping subcellular structure to their corresponding fluorescence spectra, could allow for a more comprehensive understanding of various pigmented microorganisms, which in turn will further facilitate their identification in environmental samples.

Altogether, while providing new insight into the cell biology of dinoflagellates, our study is a proof of principle for volume CLEM as a valuable tool for the ultrastructural exploration of heterogeneous environmental samples.

## Materials and methods

### Sample collection

Sampling of marine plankton was performed by towing a net of 5-10 µm mesh size for 10 minutes slowly in surface waters of the Villefranche-sur-Mer bay (France). Sampling was performed on 2021/09/14 in the early morning. Samples were filtered through serial sieves (Retsch) to collect cells measuring less than 40 µm in diameter. The fraction obtained was kept in Nalgene plastic bottles and placed in a closed cooler filled with sea water to preserve them at sea temperature and in darkness until further processing. The sample was then concentrated on a 1.2 µm size meshed mixed cellulose ester membrane (Merk) and pelleted using centrifugation for 5 minutes at 1000G and 20°C with a swinging bucket centrifuge (Eppendorf 5427R).

### High-pressure freezing (HPF) and Freeze substitution (FS)

After the collection described above, 1.2 µl of the sample pellet was loaded in a type A gold coated copper carrier (Leica microsystems, 200 µm deep and 3 mm wide) and topped with the flat side of an aluminium type B carrier (Leica microsystems). HPF was performed using an EM ICE (Leica microsystems). To allow for a very rapid freezing of the sample upon collection at sea, the instrument was set up meters away from the peer at the Institut de la mer in Villefranche-sur-Mer (France). Samples presented here were frozen within a time window of less than 2 hours after being collected at sea. Cryoimmobilized samples underwent freeze-substitution (FS, EM-AFS2, Leica microsystems) following a protocol adapted from Ronchi et al., 2021. Briefly, the samples were incubated in the FS cocktail (0.1% UA in dry acetone) for 69h at -90°C. Temperature was raised to -45°C over 15h (3°C/h) and then the samples were further incubated for 5h at -45°C. After rinsing with acetone, the infiltration with Lowicryl HM20 (Polysciences) was performed using increasing resin concentration in steps of 6h each. During infiltration the temperature was increased gradually to -25°C. Three infiltration steps using 100% Lowicryl were done at -25°C for 6h, 17h and 10h respectively. Polymerization was performed using UV at -25°C for 48h followed by raising the temperature to 20°C.

### Targeting strategy

In order to target the cell of interest we generated a 3D map of the block using confocal microscopy (Ronchi et al., 2021). For this, the sample was mounted face down on a glass bottom dish (Mattek, glass thickness 17µm) on a drop of water. Acquisition and laser branding were done using a Zeiss LSM 780 NLO microscope equipped with a pulsed near infrared (NIR) laser used in 2-photon microscopy and a 25x/0.8NA multi-immersion objective (Zeiss LD-LCI Plan-Apochromat). The following channels were acquired: two color channels detecting the autofluorescent signal of the sample, exciting autofluorescence at 488nm and 633nm respectively. Together with the 488nm excitation channel, an image of the transmitted laser light was generated using the T-PMT detector of the microscope. Additionally, a reflection channel was recorded. For this the main beamsplitter was changed to a T80/R20 filter reflecting 80% of the incident light and transmitting 20%. The reflection of a 633nm laser at low intensity was measured with a MA-PMT at low gain. Reflection protection for all laser lines was removed in the beampath of the microscope. The interface between water and resin was visible as bright reflection signal in this channel, and could be used to determine the axial position of the autofluorescent structures within the block.

For laser branding the bleaching functionality of the microscope was used, by which specific regions within an image can be selectively illuminated. For these regions the NIR laser was set to a wavelength of 850 nm. Laser power was tuned to achieve efficient branding while avoiding blebbing of the resin. With our system, we achieved this at values around 12% of the maximum power.

### Sample mounting and FIB-SEM acquisition

The block was cut parallelly to its surface in order to be 2-3 mm high, and mounted on an SEM stub (Agar scientific) using a 1:1 mix of superglue (Loctite precision max) and silver paint (EM-Tec AG44, Micro to Nano). Silver paint was further added around the block surface. The sample underwent gold sputtering for 180 s at 30 mA (Quorum Q150RS) before insertion in the FIB-SEM chamber. FIB-SEM imaging was performed using a Zeiss Crossbeam 550, following the Atlas 3D workflow. FIB milling was performed at 1.5 nA. SEM imaging was done with an acceleration voltage of 1.5 kV and a current of 750 pA using an ESB detector (ESB grid 1100V). Imaging of the planktonic cell was done using an 8 nm isotropic voxel size with a dwell time of 9 µs. Post-acquisition dataset alignment was performed using the AMST procedure from Hennies et al., 2020.

### Volume analysis and quantification

Overlay of EM and LM data (Fig. 3) and most of the segmentations (Fig. 4A,B,D,E) were done using Amira (Thermo Fisher Scientific). Additional segmentation was performed using Microscopy Image Browser (Belevich et al, 2016) (Fig. 4C). Volume quantifications were performed on segmented organelles using the Amira (Thermo Fisher Scientific) label analysis tool. Lengths of trichocysts were measured using the Amira measurement tool.

### SEM

Part of the sample collected as described above was fixed with 2% paraformaldehyde (EMS grade) and 0.5% glutaraldehyde (EMS grade) in 0.1M marPHEM (Montanaro et al., 2016) for 6h at 4°C. The sample was then transferred to 0.1M PHEM containing 1% paraformaldehyde and preserved at 4°C until further processing. The sample was then rinsed once using 0.1M PHEM at 4°C. The sample was then dehydrated at 4°C using the following (v/v) acetone/water series: 30%, 50%, 70%, 80%, 90%, followed by two pure acetone steps. Samples were left to sediment for a duration of 3 to 12h before each exchange to avoid loss of material. The sample was then critically point dried (CPD300, Leica microsystems) in small containers (1-1.6 µm pore size, Vitrapore ROBU). In the CPD program, 30 slow exchange steps were used. CPD dried plankton were then distributed on carbon tape placed on an SEM stub (Agar scientific) before further gold sputtering (Quorum Q150RS). SEM imaging was performed using a Zeiss Crossbeam 540 with an acceleration voltage of 1.5 kV and a current of 700 pA and a Secondary Electron Secondary Ion (SESI) detector.

### Datasets visualization using MoBIE

The Fiji plugin MoBIE (Pape et al., 2022) can be used to explore the different datasets. Instructions for plugin download and installation can be found using the following link: https://github.com/mobie/mobie-viewer-fiji. The data are visualized by selecting the Fiji plugin ‘MoBIE - > Open MoBIE Project’ and providing the project location (https://github.com/mobie/environmental-dinoflagellate-vCLEM). The project contains the FIB-SEM dataset (“photosynthetic dinoflagellate”) and associated segmentations, registered with the confocal stack of the full block (“LM_fullblock”), as well as the higher resolution confocal stack of the cell before and after trimming and branding (“LM_pre-trim” and “LM_trimmed-branded”).

## Data availability

The FIB-SEM datasets generated during this study are available in EMPIAR.

## Acknowledgments

We thank the EMBL and IMEV organizers of the sampling expedition in Villefranche-sur-Mer and in particular Paola Bertucci (EMBL) and Raffaela Cattaneo (IMEV, EMBR) for their support during the expedition. We thank Robert Kirmse and Leica Microsystem for their support with the HPF. We are grateful to Dr. Raffaele Siano, Dr. Hugo Bertelot, Dr. Nicolas Chomerat, Dr. Kenneth Mertens and Dr. Mona Hoppenrath for fruitful discussions and the identification of the cell species. We thank Dr Julian Hennies for help with the AMST alignment and Dr Christian Tischer for help with MoBIE. We thank Nikolaus Leisch and the Schwab team for their feedback on the manuscript. J.D., C.L., F.C. were supported by CNRS and ATIP-Avenir program. D.Y. was supported by the IDEX project of University of Grenoble Alpes (International Strategic Partnerships) and EMBL.

## Notes

### Competing Interest Statement

The authors have declared no competing interest.

## References

Belevich, I., Joensuu, M., Kumar, D., Vihinen, H. and Jokitalo, E. (2016). Microscopy Image Browser: A Platform for Segmentation and Analysis of Multidimensional Datasets. PLoS Biol. 14, 1–13.

Bhaud, Y., Guillebault, D., Lennon, J. F., Defacque, H., Soyer-Gobillard, M. O. and Moreau, H. (2000). Morphology and behaviour of dinoflagellate chromosomes during the cell cycle and mitosis. J. Cell Sci. 113, 1231–1239.

Bouck, G. B. and Sweeney, B. M. (1966). The fine structure and ontogeny of trichocysts in marine dinoflagellates. Protoplasma 61, 205–223.

Calado, A. J. and Moestrup, Ø. (2002). Ultrastructural study of the type species of Peridiniopsis, Peridiniopsis borgei (Dinophyceae), with special reference to the peduncle and flagellar apparatus. Phycologia 41, 567–584.

Calado, A. J., Hansen, G. and Moestrup, Ø. (1999). Architecture of the flagellar apparatus and related structures in the type species of peridinium, p. cinctum (dinophyceae). Eur. J. Phycol. 34, 179–191.

Colley, N. J. and Nilsson, D. E. (2016). Photoreception in Phytoplankton. Integr. Comp. Biol. 56, 764–775.

Decelle, J., Veronesi, G., LeKieffre, C., Gallet, B., Chevalier, F., Stryhanyuk, H., Marro, S., Ravanel, S., Tucoulou, R., Schieber, N., et al. (2021). Subcellular architecture and metabolic connection in the planktonic photosymbiosis between Collodaria (radiolarians) and their microalgae. Environ. Microbiol. 23, 6569–6586.

Decelle, J., Kayal, E., Bigeard, E., Gallet, B., Bougoure, J., Clode, P., Schieber, N., Templin, R., Hehenberger, E., Prensier, G., et al. (2022). Intracellular development and impact of a marine eukaryotic parasite on its zombified microalgal host. ISME J. 16, 2348–2359.

Dimprima, E., Garcia Montero, M., Gawrzak, S., Ronchi, P., Zagoriy, I., Schwab, Y., Jechlinger, M. and Mahamid, J. (2022). Integrated light and electron microscopy continuum resolution imaging of 3D cell cultures. Biophys. J. 121, 149a.

Dixon, G. K. and Syrett, P. J. (1988). The growth of dinoflagellates in laboratory cultures. New Phytol. 109, 297–302.

Dodge, J. D. (1971). Fine Structure of the Pyrrophyta. 37, 481–508.

Dodge, J. D. (1984). The functional and phylogenetic significance of dinoflagellate eyespots. BioSystems 16, 259–267.

Fensome, R. A. (1993). A classification of living and fossil dinoflagellates. Micropaleontology Press, American Museum of Natural History.

Gautier, A., Michel-Salamin, L., Tosi-Couture, E., McDowall, A. W. and Dubochet, J. (1986). Electron microscopy of the chromosomes of dinoflagellates in situ: confirmation of Bouligand’s liquid crystal hypothesis. J. Ultrastruct. Res. Mol. Struct. Res. 97, 10–30.

Gavelis, G. S., Herranz, M., Wakeman, K. C., Ripken, C., Mitarai, S., Gile, G. H., Keeling, P. J. and Leander, B. S. (2019). Dinoflagellate nucleus contains an extensive endomembrane network, the nuclear net. Sci. Rep. 9, 1–9.

Géraud, M. L., Herzog, M. and Soyer-Gobillard, M. O. (1991). Nucleolar localization of rRNA coding sequences in Prorocentrum micans Ehr. (dinomastigote, kingdom Protoctist) by in situ hybridization. BioSystems 26, 61–74.

Gomez, F. (2012). A quantitative review of the lifestyle, habitat and trophic diversity of dinoflagellates (Dinoflagellata, Alveolata). Syst. Biodivers. 10, 267–275.

Hennies, J., Lleti, J. M. S., Schieber, N. L., Templin, R. M., Steyer, A. M. and Schwab, Y. (2020). AMST: Alignment to Median Smoothed Template for Focused Ion Beam Scanning Electron Microscopy Image Stacks. Sci. Rep. 10, 1–10.

Hense, B. A., Gais, P., Jütting, U., Scherb, H. and Rodenacker, K. (2008). Use of fluorescence information for automated phytoplankton investigation by image analysis. J. Plankton Res. 30, 587–606.

Hoppenrath, M. (2017). Dinoflagellate taxonomy — a review and proposal of a revised classification. Mar. Biodivers. 47, 381–403.

Introduction to Electron Microscopy of Cells (2007). Methods Cell Biol. 79,.

Kukulski, W., Schorb, M., Welsch, S., Picco, A., Kaksonen, M. and Briggs, J. A. G. (2011). Correlated fluorescence and 3D electron microscopy with high sensitivity and spatial precision. J. Cell Biol. 192, 111–119.

Li, Z., Mertens, K. N., Gottschling, M., Gu, H., Söhner, S., Price, A. M., Marret, F., Pospelova, V., Smith, K. F., Carbonell-Moore, C., et al. (2020a). Taxonomy and Molecular Phylogenetics of Ensiculiferaceae, fam. nov. (Peridiniales, Dinophyceae), with Consideration of their Life-history. Protist 171,.

Li, Z., Mertens, K. N., Gottschling, M., Gu, H., Söhner, S., Price, A. M., Marret, F., Pospelova, V., Smith, K. F., Carbonell-Moore, C., et al. (2020b). Taxonomy and Molecular Phylogenetics of Ensiculiferaceae, fam. nov. (Peridiniales, Dinophyceae), with Consideration of their Life-history. Protist 171, 125759.

Moestrup, √òjvind and Daugbjerg, N. (2007). On dinoflagellate phylogeny and classification. 215–230.

Moldrup, M., Moestrup, Ø. and Hansen, P. J. (2013). Loss of phototaxis and degeneration of an eyespot in long-term algal cultures: Evidence from ultrastructure and behaviour in the dinoflagellate Kryptoperidinium foliaceum. J. Eukaryot. Microbiol. 60, 327–334.

Montanaro, J., Gruber, D. and Leisch, N. (2016). Improved ultrastructure of marine invertebrates using non-toxic buffers. PeerJ 2016, 1–15.

Nixon, S. J., Webb, R. I., Floetenmeyer, M., Schieber, N., Lo, H. P. and Parton, R. G. (2009). A single method for cryofixation and correlative light, electron microscopy and tomography of zebrafish embryos. Traffic 10, 131–136.

Oliveira, C. Y. B., Oliveira, C. D. L., Müller, M. N., Santos, E. P., Dantas, D. M. M. and Gálvez, A. O. (2020). A Scientometric Overview of Global Dinoflagellate Research. Publications 8, 50.

Pape, C., Meechan, K., Moreva, E., Schorb, M., Chiaruttini, N., Zinchenko, V., Vergara, H., Mizzon, G., Moore, J., Arendt, D., et al. (2022). MoBIE: A Fiji plugin for sharing and exploration of multi-modal cloud-hosted big image data. bioRxiv 2022.05.27.493763.

Peddie, C. J., Genoud, C., Kreshuk, A., Meechan, K., Micheva, K. D., Narayan, K., Pape, C., Parton, R. G., Schieber, N. L., Schwab, Y., et al. (2022). Volume electron microscopy. Nat. Rev. Methods Prim. 2, 51.

Plattner, H. (2017). Trichocysts—Paramecium’s Projectile-like Secretory Organelles: Reappraisal of their Biogenesis, Composition, Intracellular Transport, and Possible Functions. J. Eukaryot. Microbiol. 64, 106–133.

Porrati, F., Grewe, D., Seybert, A., Frangakis, A. S. and Eltsov, M. (2019). FIB-SEM imaging properties of Drosophila melanogaster tissues embedded in Lowicryl HM20. J. Microsc. 273, 91–104.

Ronchi, P., Mizzon, G., Machado, P., D’imprima, E., Best, B. T., Cassella, L., Schnorrenberg, S., Montero, M. G., Jechlinger, M., Ephrussi, A., et al. (2021). High-precision targeting workflow for volume electron microscopy. J. Cell Biol. 220,.

Seo, K. S. and Fritz, L. (2002). Diel changes in pyrenoid and starch reserves in dinoflagellates. Phycologia 41, 22–28.

Truby, E. W. (1997). Preparation of single-celled marine dinoflagellates for electron microscopy. Microsc. Res. Tech. 36, 337–340.

Uwizeye, C., Brisbin, M. M., Gallet, B., Chevalier, F., LeKieffre, C., Schieber, N. L., Falconet, D., Wangpraseurt, D., Schertel, L., Stryhanyuk, H., et al. (2021a). Cytoklepty in the plankton: A host strategy to optimize the bioenergetic machinery of endosymbiotic algae. Proc. Natl. Acad. Sci. U. S. A. 118,.

Uwizeye, C., Decelle, J., Jouneau, P.-H., Flori, S., Gallet, B., Keck, J.-B., Bo, D. D., Moriscot, C., Seydoux, C., Chevalier, F., et al. (2021b). Morphological bases of phytoplankton energy management and physiological responses unveiled by 3D subcellular imaging. Nat. Commun. 12, 1049.

Vargas, C., Audic, S., Henry, N., Decelle, J., Mahé, F., Logares, R., Lara, E., Berney, C., Le Bescot, N., Probert, I., et al. (2015). Eukaryotic plankton diversity in the sunlit ocean. Science (80-.). 348, 1261605–1/11.

Westermann, M., Steiniger, F., Gülzow, N., Hillebrand, H. and Rhiel, E. (2015). Isolation and characterisation of the trichocysts of the dinophyte Prorocentrum micans. Protoplasma 252, 271–281.

