## Supplementary figures and legends for "Targeted volume Correlative Light and Electron Microscopy of an environmental marine microorganism"

### Supplementary material

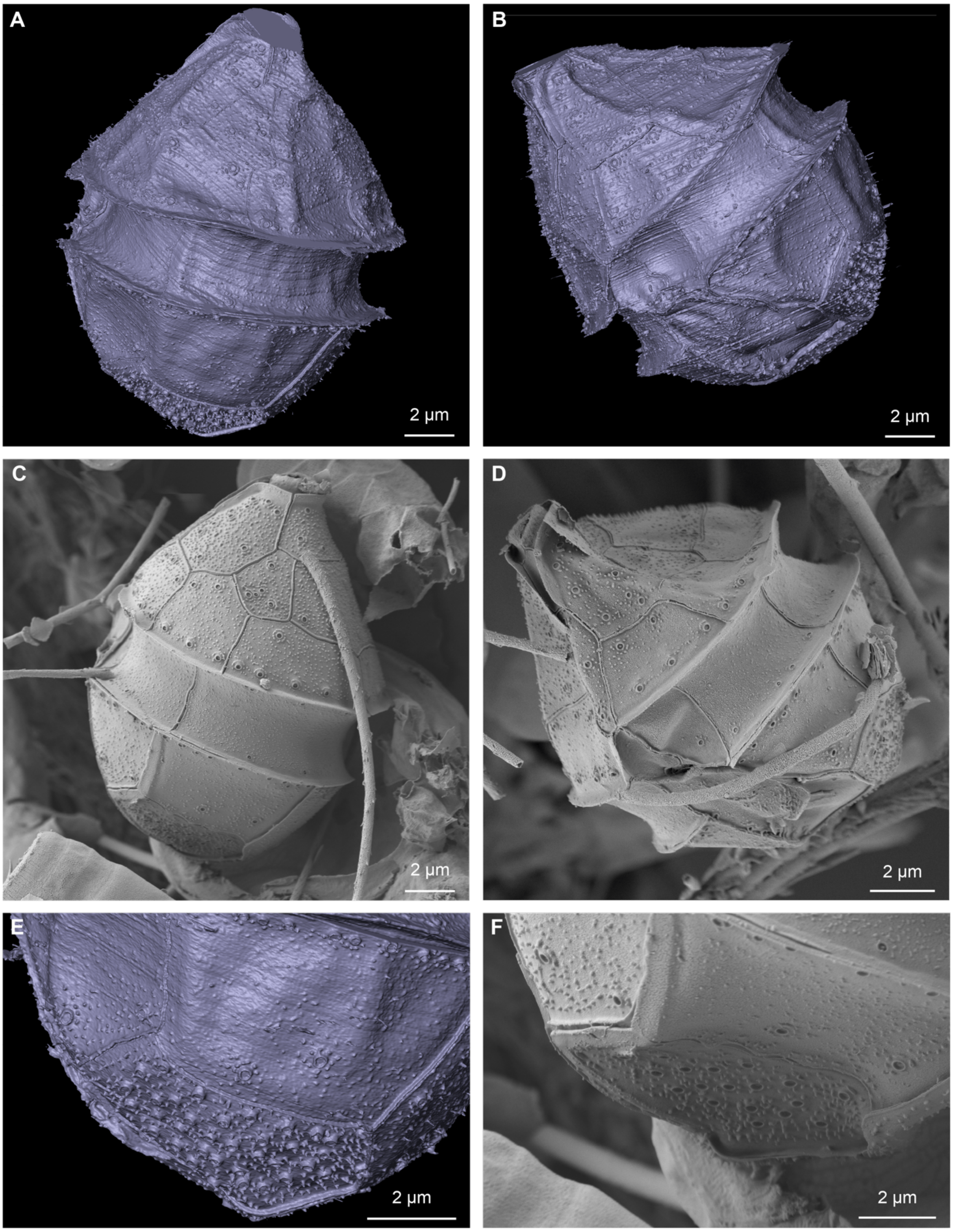


**Figure S1. Theca shape and ornamentation as visualized from vEM data compared to topography SEM analysis of the same species.**
A,B) 3D rendering of the theca of the cell, as segmented from FIB-SEM data.
C,D) Topography SEM images of cells collected in parallel to the ones prepared for FIB-SEM. Images show cells that we believe correspond to the same species (*Ensiculifera*

*tyrrhenica),* captured in the same orientation as the views of the rendering in A and B, respectively.

E) Rendering of the FIB-SEM data. Detailed view of adjacent thecal plates, showing the spatial organization of the openings and knobs.

F) Higher magnification view of the cell shown in C, visualizing the ornamentation of the thecal plates, matching to E.

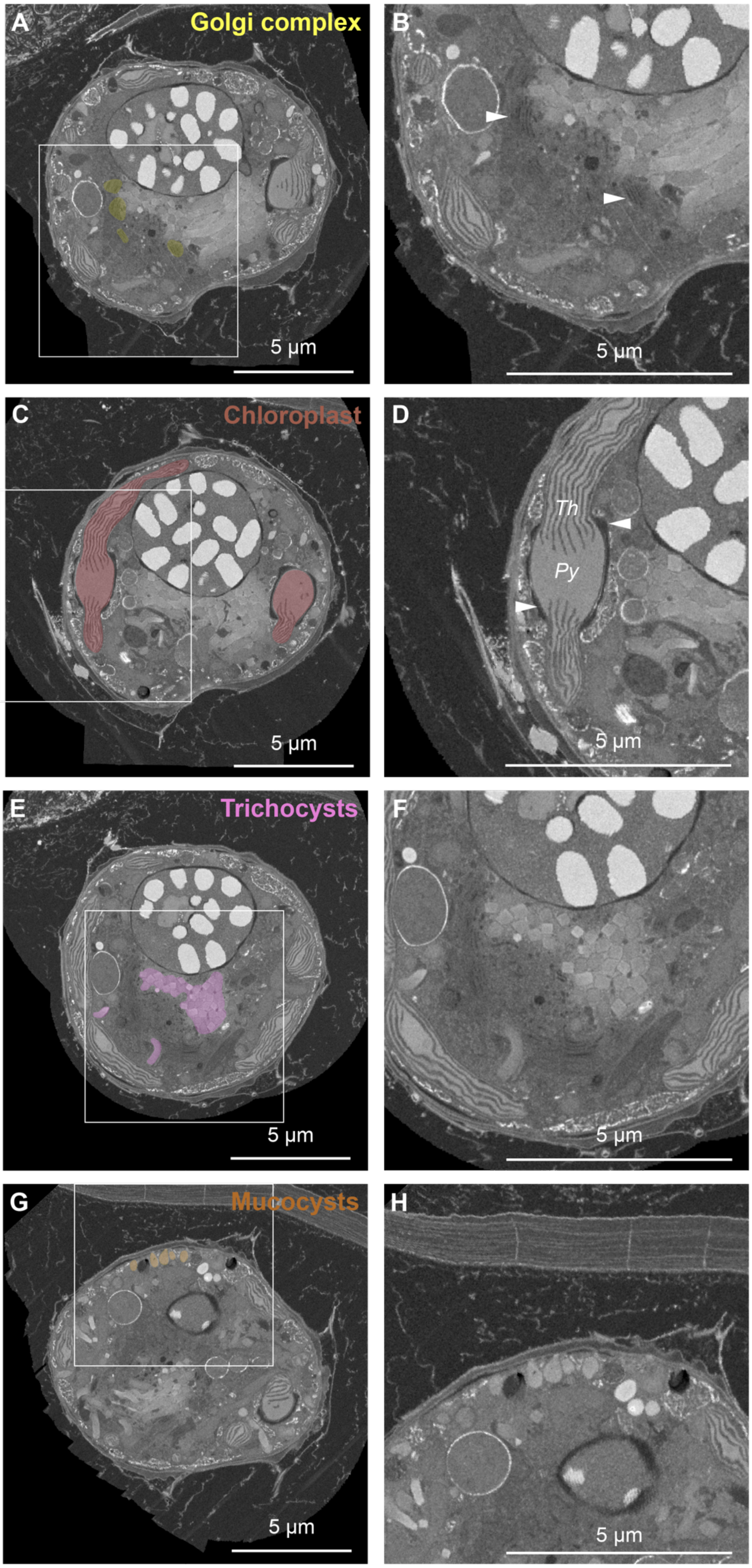
A
Golgi complex
5 µm
B
5 µm
C
Chloroplast
5 µm
D
Th
Py
5 µm
E
Trichocysts
5 µm
F
5 µm
G
Mucocysts
5 µm
H
5 µm

**Figure S2. Raw images from the FIB-SEM volume showing ultrastructural details of the different organelles.**
A low magnification image with segmentation overlay is shown for each organelle in the left panels. The boxed areas are magnified in the right panel.
A,B) Golgi complex. In B, arrowheads highlight two Golgi stacks with characteristic cisternae organization.
C,D) Chloroplast. In D, the organization of the pyrenoid (Py) and thylacoid membranes (Th) is shown. Arrowheads point at the starch accumulation around the chloroplast in the pyrenoid region.
E,F) Trychocysts. F shows the characteristic cross section squared profiles.
G,H) Mucocysts. The typical amphora shape and sub-plasma membrane location are hallmarks of these secretory organelles.

**Video S1. vCLEM targeting of a photosynthetic dinoflagellate in an heterogeneous sample.**
The video shows the progressive targeting of the cell of interest from the confocal stack of a resin block to EM imaging, followed by overlay of the volumes acquired in the two modalities. The video has been generated from the MoBIE project.

**Video S2. Segmentation of the FIB-SEM dataset.**
The video shows orthoslices through the volume and rendering of some representative structures. The theca is shown in grey, flagella in green, nuclear envelope in cyan, Golgi apparatus in yellow, chloroplast in red, with the associated starch in white, chromosomes in blue, trichocysts in pink and purple (short and long class, respectively), mucocysts in orange.
